# An efficient FLP-based toolkit for spatiotemporal control of gene expression in *Caenorhabditis elegans*

**DOI:** 10.1101/107029

**Authors:** Celia María Muñoz-Jiménez, Cristina Ayuso, Agnieszka Dobrzynska, Antonio Torres, Patricia de la Cruz Ruiz, Peter Askjaer

**Affiliations:** Andalusian Center for Developmental Biology (CABD), CSIC/JA/Universidad Pablo de Olavide, 41013 Seville, Spain.

**Keywords:** *baf-1*, DamID, FLP-out gene inactivation, genome engineering, MosSCI, PEEL-1 toxin, tissue-specific gene expression.

## Abstract

Site-specific recombinases are potent tools to regulate gene expression. In particular, the Cre and FLP enzymes are widely used to either activate or inactivate genes in a precise spatiotemporal manner. Both recombinases work efficiently in the popular model organism *Caenorhabditis elegans* but their use in this nematode is still only sporadic. To increase the utility of the FLP system in *C. elegans* we have generated a series of single-copy transgenic strains that stably express an optimized version of FLP in specific tissues or by heat induction. We show that recombination efficiencies reach 100 percent in several cell types, such as muscles, intestine and serotonin producing neurons. Moreover, we demonstrate that most promoters drive recombination exclusively in the expected tissues. As examples of the potentials of the FLP lines we describe novel tools for induced cell ablation by expression of the PEEL-1 toxin and a versatile FLP-out cassette for generation of GFP-tagged conditional knockout alleles. Together with other recombinase-based reagents created by the *C. elegans* community this toolkit increases the possibilities for detailed analyses of specific biological processes at developmental stages inside intact animals.

## Introduction

Precise manipulation of gene expression is central to understand most biological processes. Strategies such as gene knockouts, expression of transgenes and RNA-mediated interference are commonly used tools but many experiments also require spatiotemporal regulation; that is either activation or inactivation of (trans-) genes in specific cell types and/or at particular moments during the experiment. Efficient optogenetics methods have emerged to achieve this, but they require certain technical expertise and are most suitable to analyze a relatively low number of cells and animals (Qi *et al.* 2012; Churgin *et al.* 2014; Glock *et al.* 2015; Xu and Chisholm 2016). Genetic approaches based on site-specific recombinases, such as FLP (flipase) and Cre (cyclization recombination) derived from the yeast *Saccharomyces cerevisiae* and the bacteriophage P1, respectively, have been adapted to induce DNA excisions, inversions or translocations in a variety of organisms, including several plants (Liu *et al.* 2013), mouse (Anastassiadis *et al.* 2010), fish (Trinh le *et al.* 2011; Wong *et al.* 2011), fruit fly (Xu and Rubin 2012) and nematodes (Hubbard 2014). FLP and Cre recognize a specific 34-bp recombination target (RT) sequence known as Frt (FLP recognition target) and loxP (locus of crossing over in P1), respectively. The RT sites consist of a central, non-palindromic 8-bp sequence flanked by 13-bp inverted perfect (wild type loxP) or imperfect (Frt) repeats. Two molecules of the recombinase bind cooperatively to an RT site and if two compatible RTs are within the vicinity of each other recombination will occur between the central 8-bp sequences. If the two RT sites are positioned on the same DNA molecule and in the same orientation, the recombination reaction will excise the intervening sequence and leave a single RT site (Figure 1). If the RTs are oriented towards each other, the intervening sequence will be inverted, whereas reactions involving two pairs of RT sites can be used for recombinase-mediated cassette exchange. Normally, the recombination events catalyzed by FLP and Cre are reversible but variation in Frt and loxP sequences have been explored to make reactions irreversible and to increase the control of multiple recombination events between homotypic and heterotypic recognition sites (Anastassiadis *et al.* 2010; Hubbard 2014).

**Figure 1.**
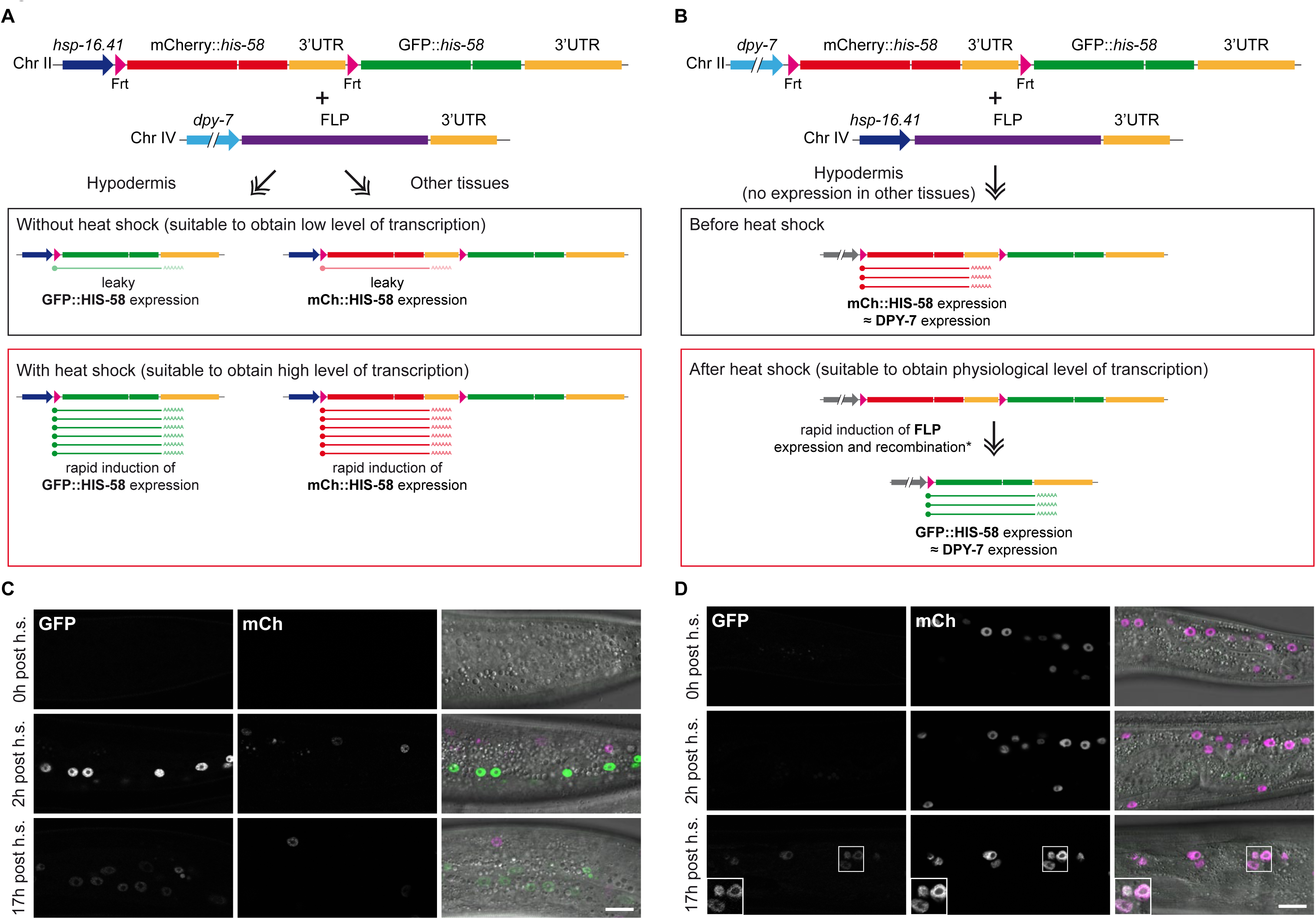
Comparison of strategies for spatiotemporal control of gene expression. (AB) Schematic representation of dual color reporters integrated by MosSCI into chromosome II, and constructs expressing FLP recombinase inserted into chromosome IV. The dual color reporters contain a Frt-flanked mCherry::*his-58* cassette with a *let-858* 3’UTR and a downstream GFP::*his-58* cassette with a *unc-54* 3’UTR, whereas the *glh-2* 3’UTR is used in the FLP constructs. (A) Placing the FLP and Frt transgenes under control of a tissue-specific promoter (*dpy-7* is used as example) and the heat-inducible *hsp-16.41* promoter, respectively, enables very rapid, strong and transient induction of a downstream transgene (GFP::*his-58* in this example) is the target tissue. In other tissues, induction of the *hsp-16.41* promoter leads to expression of mCherry::HIS-58. Because of the leaky expression of the *hsp-16.41 promoter* in the absence of induction, the system can be utilized to obtain low expression in the target tissue. (B) In the opposite configuration, in the absence of heat-induction, the Frt-flanked transgene is expressed in the target tissue at levels equivalent to the activity of the tissue-specific promoter. A heat shock will lead to high expression of FLP in all tissues and efficient recombination. After recombination (i.e. after a short time delay indicated by an asterisk), the downstream transgene is under control of the tissue-specific promoter. In A, the target tissue includes all cells derived from lineages in which the tissue-specific promoter has induced recombination; in B, the downstream gene will only be expressed when and where the tissue-specific promoter is active. (C-D) Single confocal sections of animals carrying the transgenes described in A (C) and B (D). Shown are examples 0 h (top), 2 h (middle) and 17 h (bottom) after heat induction. Note the rapid and transient induction of GFP::HIS-58 (green in merge; mCherry in magenta) in C, which is in contrast to the slow induction in D. Images were acquired with identical microscope settings; inserts in D represent contrast adjusted images to facilitate visualization of GFP::HIS-58. Scale bar 10 μm.

There are multiple considerations for optimal design of a recombination system. FLP and Cre both work efficiently in mammalian cells, and particularly the latter has gained popularity in the design of transgenic mice although high expression levels of Cre can be toxic to mammalian cells (see (Anastassiadis *et al.* 2010; Takata *et al.* 2011) for references). The original FLP enzyme has optimal activity at 30°C, whereas recombination by Cre is highest around 37°C (Buchholz *et al.* 1996). This might have favored the creation of Cre-related tools in mammalian systems although FLP variants with increased temperature stability have been developed (Buchholz *et al.* 1998). A direct comparison of FLP and Cre activities suggests that the optimal recombinase for a particular reaction might differ between recombination strategies (Takata *et al.* 2011). The combined use of Frt and loxP sites in allele construction can provide additional layers of control of gene expression (Anastassiadis *et al.* 2010).

Whereas in most organisms the use of recombinases to control gene expression involves stable lines where FLP or Cre expression constructs are inserted into the genome and crossed to lines carrying Frt or loxP-containing transgenes, experiments in *Caenorhabditis elegans* have generally been performed with expression of recombinases from extrachromosomal arrays or injected mRNA (Hubbard 2014). Extrachromosomal arrays are easy to generate in *C. elegans* and their variable stability during mitotic divisions can be explored through analyses of mosaic animals. Nevertheless, for many experiments, it is desirable that recombination occurs in most or all cells of a particular tissue and in a reproducible manner between individual animals in a population. The Hubbard and Jorgensen laboratories were the first to report the utility of the FLP/Frt system in *C. elegans* (Davis *et al.* 2008; Voutev and Hubbard 2008). Both studies found that FLP expressed from extrachromosomal arrays was able to excise (“flip out”) transgenes that were also present in multiple copies. For concatenated multi-copy Frt reporter constructs different outcomes can be expected, ranging from no excision to partial excision (i.e. some but not all Frt-flanked cassettes are flipped out from the multi-copy transgene) and complete excision. Generally, partial excision was observed in the aforementioned studies although some cells underwent complete excision (Davis *et al.* 2008; Voutev and Hubbard 2008). Voutev and Hubbard also quantified FLP activity in a single cell by expressing FLP from integrated constructs carrying the *hsp-16.2* heat shock promoter and observed recombination of an extrachromosomal reporter in the dorsal cell of gonadal sheath pair 1 in 11-20% of the animals.

Possibly, the complexity of the multi-copy reporter was partly responsible for this relatively low efficiency. Importantly, expression of FLP was reported not to affect viability, fecundity or behavior (Voutev and Hubbard 2008).

Direct comparison of FLP and Cre in the same cell types and on identical reporters is still limited in *C. elegans*. Intersectional restriction of gene expression involves two different cell type-specific promoters to drive expression of the recombinase and its target construct: the gene downstream of the RT cassette is expressed only in tissues where both promoters are active. A recent study tested the FLP and Cre systems to achieve intersectional restriction of expression and concluded that it depends on the promoter combinations which recombinase works best (Schmitt *et al.* 2012). Remarkably, highly restricted expression in one or two neurons with either FLP/Frt or Cre/loxP can be achieved in close to 100% of tested animals although extensive optimization might be required (Macosko *et al.* 2009; Ezcurra *et al.* 2011; Schmitt *et al.* 2012). Insertion of Frt or loxP sites into endogenous loci have also proven to be feasible for either complete gene knockout (Vazquez-Manrique *et al.* 2010) or conditional deletion of a genomic ~1 kb fragment (Lo *et al.* 2013). In addition, Cre has successfully been implemented for removal of selection markers after CRISPR-mediated genome engineering (Dickinson and Goldstein 2016; Schwartz and Jorgensen 2016) as well as for control of loxP-flanked integrated single-copy transgenes (Flavell *et al.* 2013; Ruijtenberg and van den Heuvel 2015).

To facilitate the use of the FLP/Frt recombination system to manipulate gene expression in a highly efficient, versatile and reproducible manner, we decided to generate a collection of stable FLP lines. We provide a characterization of 13 integrated, single-copy FLP lines that cover all major tissue types in *C. elegans* and report their efficiencies at single-cell resolution. We demonstrate that the system offers tight spatiotemporal control of expression of a fluorescent reporter protein and as further examples of their utility we developed tools for conditional cell ablation and knockout of CRISPR-engineered GFP-tagged alleles. Importantly, the tools presented here are compatible with other reagents created by the *C. elegans* community and together they increase the potentials of the gene expression toolbox.

## Materials and Methods

### Plasmid construction

We initially PCR-amplified a FLP::3’UTR*_glh-2_* sequence from the plasmid pWD79 (Davis *et al.* 2008) with primers B441 and B551 (Supporting Information, Table S1) and inserted it into *Kpn*I/*Xma*I of plasmid pBN9 designed for Mos1-mediated single copy insertion (MosSCI) into chromosome IV (Morales-Martinez *et al.* 2015). This FLP version, which we here name FLP pWD79, encodes a glycine residue at position 5 and contains a single artificial intron (see Supporting Information for complete sequences of FLP versions used in this study). Next, we inserted a 983 bp *Pst*I/*Kpn*I P*_myo-2_* fragment from pBN41 (Morales-Martinez *et al.* 2015) upstream of the FLP pWD79 sequence to generate plasmid pBN155 P*_myo-2_*::FLP pWD79. Plasmids pBN158 P*_myo-3_*::FLP pWD79, pBN159 P*_unc-119_*::FLP pWD79 and pBN160 P*_eft-3_*::FLP pWD79 were made by substituting the *Pst*I/*Kpn*I P*_myo-2_* sequence of pBN155 by the corresponding promoter sequences amplified from Bristol N2 genomic DNA (or plasmid pUP1 (Rodenas *et al.* 2012) for *unc-119*) using a KAPA HiFi DNA polymerase (KAPA Biosystems, Wilmington, MA). Throughout this study we used conventional restriction enzyme-mediated cloning and Gibson assembly technology. By introducing relatively few modifications, our plasmids can be made Gateway compatible but others have observed that the extra sequences inherent to Gateway recombination sites might decrease recombination efficiencies (Schmitt *et al.* 2012). The primers used to clone promoters in this study as well as promoter lengths and sequence verification are described in Table S2 in the Supporting Information.

To increase FLP expression levels we used the *C. elegans* codon adapter tool (Redemann *et al.* 2011) with the parameters “Optimize for weak mRNA structure at ribosome binding site” and “Avoid splice sites in coding region” resulting in a codon adaptation index (CAI) of 0.9808. The sequence, which we termed FLP G5, was synthesized by Eurofins Genomics (Ebersberg, Germany). The initial design contained 3 artificial introns but due to cloning problems the final sequence contains 2 introns as well as the *glh-2* 3’UTR. The FLP G5::3’UTR*_glh_*_-*2*_ sequence is flanked by *Kpn*I and *Xma*I sites and was used to substitute the *Kpn*I/*Xma*I fragment of pBN155 to create plasmid pBN168 P*_myo-2_*::FLP G5. Derivatives of pBN168 were made by substituting the *myo-2* promoter by other promoters as either *Pst*I/*Kpn*I or *Not*I/*Kpn*I fragments from PCR amplicons or plasmids to generate pBN172 P*_myo-3_*::FLP G5, pBN173 P_*unc-119*_::FLP G5, pBN176 P*_nhr-82_*::FLP G5 and pBN219 P*_dpy-7_*::FLP G5.

Substitution of glycine for aspartic acid at position 5 to generate FLP D5 was made by PCR amplification using FLP G5 as template and primers B572 and B826. The forward primer introduced two nucleotide changes in the fifth FLP codon and the resulting fragment was digested with *Kpn*I and *Sal*I and used to replace the *Kpn*I/*Sal*I fragment of pBN176 to generate pBN237 P*_nhr-82_*::FLP D5. Derivatives of pBN237 were made by substituting the *nhr-82* promoter by other promoters by either conventional cloning of *Not*I/*Kpn*I fragments (from PCR amplicons or subcloning) or Gibson assembly to generate pBN258 P*_unc-119_*::FLP D5, pBN260 P*_myo-3_*::FLP D5, pBN262 _P*hsp-16.41*_::FLP D5, pBN263 P*_myo-2_*::FLP D5, pBN266 P*_dpy-7_*::FLP D5, pBN267 P*_rgef-1_*::FLP D5, pBN278 P*_mec-7_*::FLP D5, pBN279 P*_hlh-8_*::FLP D5, pBN280 P*_tph-1_*::FLP D5, pBN282 P*_elt-2_*::FLP D5, pBN283 P*_dat-1_*::FLP D5, pBN303 P*_unc-47_*::FLP D5 and pBN310 P*_lag-2_*::FLP D5. Sequence verification revealed that pBN282 carries a mutation that introduces an amino acid substitution at position 6 of FLP D5 (I6L), which does not prevent enzymatic activity.

Finally, FLAG-NLS-FLP D5 was made by Gibson assembly using a 222 nt gBlock (Integrate DNA Technologies, Leuven, Belgium) encoding 3xFLAG and a nuclear localization sequence (NLS) from EGL-13, a PCR fragment encoding FLP D5 (primers B827+B828), a PCR fragment containing the *tbb-2* 3’UTR (primers B829+B830) and pBN176 digested with *Kpn*I and *Xma*I to generate pBN238 P*_nhr-82_*::FLAG-NLS-FLP D5. These modifications were inspired by results with the Cre recombinase (Ruijtenberg and van den Heuvel 2015). Derivatives of pBN238 were made by substituting the *nhr-82* promoter by other promoters by either conventional cloning of *Not*I/*Kpn*I fragments from plasmids described above or Gibson assembly to generate pBN252 P*_dat-1_*::FLAG-NLS-FLP D5, pBN254 P*_unc-47_*::FLAG-NLS-FLP D5, pBN261 P*_hsp-16.41_*::FLAG-NLS-FLP D5, pBN268 P*_rgef-1_*::FLAG-NLS-FLP D5, pBN269 P*_unc-119_*::FLAG-NLS-FLP D5 and pBN270 P*_myo-3_*::FLAG-NLS-FLP D5.

Plasmid pBN338 P*_dat-1_*::FLP D5::SL2::mNG was designed to provide visualization of FLP D5 expression. First, an intergenic *gpd-2/-3* fragment and a mNeonGreen sequence were amplified with primers B1012+B1013 and B1014+B1015 using pMLS268 (Schwartz and Jorgensen 2016) and dg357 (Hostettler *et al.* 2017) as templates, respectively. Next, the two products were stitched by PCR with primers B1012 and B1015 and inserted by Gibson assembly into *Mlu*I of pBN283 followed by sequence verification.

Whereas the FLP expression plasmids described here all are designed for MosSCI integration into chromosome IV, most promoter::FLP::3’UTR cassettes (including cassettes with *dat-1*, *eft-3*, *myo-2*, *nhr-82*, *rgef-1*, *unc-47* or *unc-119* promoters) can be excised by digestion with *Not*I and *Xma*I and inserted into *Not*I/*Ngo*MIV of pBN8 (Rodenas *et al.* 2012). Plasmid pBN8 was originally designed for insertion into chromosome II by MosSCI but can be used for insertion into any of the universal MosSCI sites generated on chromosomes I-V (see http://www.wormbuilder.org/ and (Frokjaer-Jensen *et al.* 2014).

The heat inducible dual color reporter plasmid pBN154 P*_hsp-16.41_*>mCherry::*his-58*::3’UTR*_let_*_-*858*_>gfp::*his-58*::3’UTR*_unc_*_-*54*_ (Frt recombination sites are indicated with “>” symbols) was generated through multiple steps. 1) A >mCherry::3’UTR*_let-858_*>gfp cassette was amplified from pWD178 (Davis *et al.* 2008) with primers B445 and B446 and inserted into the cloning vector pSPARK I (Canvax Biotech, Cordoba, Spain) to produce plasmid #1151. 2) Plasmid #1151 was digested with *Not*I and religated to produce plasmid #1231. 3) The >mCherry::3’UTR*_let-858_*>gfp cassette was purified from plasmid #1231 by digestion with *Sal*I and *Xma*I and inserted into *Xho*I/*Ngo*MIV of pBN16 (Rodenas *et al.* 2012) to generate plasmid pBN152. 4) A *his-58*-containing *Xho*I/*Bam*HI fragment from plasmid pJH4.52 (Strome *et al.* 2001) was inserted into *Xho*I/*Bgl*II of pBN152 to produce pBN153. 5) Two overlapping PCR fragments containing P*_hsp-16.41_*>mCherry 5’half (amplified with primers B211 and B440 using pBN153 as template) and mCherry 3’half::*his-58* (amplified with primers B220 and B355 using pBN1 (Rodenas *et al.* 2012) as template) were PCR-stitched with primers B211 and B355 and inserted into pSPARK I. 6) Finally, a *Sph*I/*Stu*I fragment encoding P*_hsp-16.41_*>mCherry::*his-58* from this plasmid was inserted into *Sph*I/*Nae*I of pBN153 to generate pBN154. All inserts were sequence verified. The hypodermal dual color reporter plasmid pBN368 P*_dpy-7_*>mCherry::*his-58*::3’UTR*_let-858_*>gfp::*his58*::3’UTR*_unc_*_-*54*_ was generated in two steps. 1) The >mCherry::*his-58*::3’UTR*_let-858_*>gfp::*his-58*::3’UT*_Runc_*_-*54*_ sequence was amplified from pBN154 with primers B1064 and B1065 and inserted by Gibson assembly into pBN266 cut with *Kpn*I and *Xma*I to generate pBN360. 2) A 2.5 kb *Sph*I/*Xho*I fragment from pBN360 was inserted into *Sph*I/*Xho*I of pBN154 to produce pBN368.

Plasmid pBN189 P*_hsp-16.41_*>mCherry::*his-58*::3’UTR*_let-858_*>Dam::*mel-28*::3’UTR*_unc_*_-*54*_ for FLP-induced expression of Dam::MEL-28 was generated by Gibson assembly using two overlapping PCR fragments containing P*_hsp-16.41_*>mCherry::*his-58*::3’UTR*_let-858_*> (amplified with primers B662 and B676 using pBN154 as template) and Dam (amplified with primers B674 and B675 using pBN69 P*_hsp_*_-*16.41*_::Dam::*mel-28*::3’UTR*_unc_*_-*54*_ (Sharma *et al.* 2014) as template) together with pBN69 digested with *Not*I and *Ngo*MIV. Next, pBN189 was digested with *Ngo*MIV and *Nhe*I to substitute the *mel-28* sequence by a *Ngo*MIV/*Nhe*I fragment from pBN79 P*_hsp-16.41_*::Dam::*emr-1*::3’UTR*_unc-54_* (Gonzalez-Aguilera *et al.* 2014) to produce pBN209 P*_hsp-16.41_*>mCherry::*his-58*::3’UTR*_let-858_*>Dam::emr-1::3’UTR*_unc-54_*. Finally, pBN209 was digested with *Xho*I and *Nhe*I to substitute the Dam::*emr-1* sequence by a *Xho*I/*Nhe*I *peel-1* cDNA fragment amplified by PCR with primers B871 and B872 and pMA122 (Frokjaer-Jensen *et al.* 2012) as template, producing plasmid pBN255 P*_hsp-16.41_*>mCherry::*his-58*::3’UTR*_let-858_*>*peel-1*::3’UTR*_unc-54_*. All inserts were sequence verified.

Plasmid pBN312 GFP KO for FLP-mediated gene knockout was generated by insertion of two synthetic DNA fragments (IDT, Leuven Belgium) into *Nco*I+*Xba*I of pMLS252 (a gift from Erik Jorgensen; Addgene plasmid # 73720). pBN312 contains 34 bp Frt sequences in the first and second intron of GFP as well as an *unc-119* rescuing fragment flanked by loxP sites in the third GFP intron. pBN312 can be used as “Tag + Marker Donor Plasmid” in the recent SapTrap protocol for CRISPR/Cas9 genome engineering (Schwartz and Jorgensen 2016). The targeting vector pBN313 for insertion of the GFP knockout cassette into the *baf-1* locus was made by SapTrap cloning with destination plasmid pMLS256, N-tagging connector plasmid pMLS288 and tag + donor plasmid pBN312 as well as the annealed primer pairs 5'HA (B963- B964), 3'HA (B965-B966) and sg (B961-B962). All inserts were sequence verified.

Plasmid pBN306 P*_myo_*_-*2*_::GFP::3’UTR*_unc_*_-*54*_ was generated by subcloning a *Pst*I/*Spe*I fragment from pBN41 into *Pst*I/*Spe*I of pBN9. Most plasmids produced in this study and their sequences are available at Addgene (www.addgene.org); remaining plasmids can be requested from the authors.

### Strains and transgenesis

Strains were maintained on standard NGM plates at 16°C (Stiernagle 2006). Except where noted, all transgenic strains were made by Mos1-mediated single copy insertion (MosSCI) with *unc-119(+)* recombination plasmids together with pCFJ601 (transposase) and three red (pCFJ90, pCFJ104 and pBN1) or green (pBN40, pBN41 and pBN42) co-injection markers (Frokjaer-Jensen *et al.* 2012; Dobrzynska *et al.* 2016). Plasmids were microinjected into the gonads of uncoordinated adults and successful integrants were identified after 7-14 days based on wildtype locomotion and absence of fluorescent markers. FLP transgenes were inserted into the *cxTi10882* locus in chromosome IV of strain EG5003 *unc-119(ed3)* III or EG6700 *unc-119(ed3)* III. Frt constructs were inserted into the *ttTi5605* locus in chromosome II of strain EG4322 *unc-119(ed9)* III. Strains BN225 (Morales-Martinez *et al.* 2015), BN243 (Gonzalez-Aguilera *et al.* 2014) and BN578 carrying fluorescent markers inserted into the *cxTi10882* and *ttTi5605* loci were used to facilitate crosses. Strains OH7193 (Flames and Hobert 2009) and XE1375 (Firnhaber and Hammarlund 2013) were used to mark dopaminergic and GABAergic neurons, respectively.

Strain BN552, which carries a GFP knockout cassette and an *unc-119* rescuing gene inserted into the *baf-1* locus, was generated by injection of HT1593 *unc-119(ed3)* with plasmids pBN313 (65 ng/μl), #1286 P*_eft-3_*::cas9-SV40-NLS (25 ng/μl; (Friedland *et al.* 2013)), pBN1 (10 ng/μl), pCFJ90 (2.5 ng/μl) and pCFJ104 (5 ng/μl) and selecting for non-red, wildtype-moving animals. Next, the *unc-119* rescuing gene in the third intron of GFP in BN552 was removed to generate BN565. This was achieved by injection with Cre plasmid pMLS328 (50ng/ul; (Schwartz and Jorgensen 2016)), pBN1 (10 ng/μl), pCFJ90 (2.5 ng/μl), pCFJ104 (5 ng/μl) and an irrelevant plasmid to reach a total DNA concentration of 100 ng/μl, followed by selection for non-red, Unc animals. Finally, BN565 was crossed with wildtype N2 to produce BN580, which is *unc-119(+)*, and with BN508 P*elt-2*:: FLP D5 to produce BN582, in which *baf-1* expression is disrupted specifically in intestinal cells.

See Supporting Information, Table S3 for complete list of strains. At least one strain for each FLP D5 line will be made immediately available through the Caenorhabditis Genetics Center; others can be requested from the authors.

### Microscopy

Evaluation of FLP activity on the dual color reporter was assayed by heat induction at 34°C for 15 min followed by recovery at 20°C for ~3 h unless indicated otherwise. In experiments with two heat shocks, these were performed 24 h apart. For confocal microscopy, nematodes were mounted in 3 μL 10 mM levamisole HCl (Sigma-Aldrich, St. Louis, MI, USA) on 2% agarose pads (Askjaer *et al.* 2014) and observed on a Nikon Eclipse Ti microscope equipped with Plan Fluor 40x/1.3 and Plan Apo VC 60x/1.4 objectives and a A1R scanner (Nikon, Tokyo, Japan) using a pinhole of 1.2–1.4 airy unit. Quantification of GFP positive nuclei was done by direct observation at the microscope, or, in most cases, after acquisition of z-stack images with integrated Nikon NIS software. Image stacks were opened in Fiji/ImageJ 2.0.0-rc-43 (Schindelin *et al.* 2012), which was also used for post-acquisition processing, such as cropping, merging and maximum projection.

### peel-1-induced toxicity

For strains with FLP D5 expressed in body wall muscles (P*_myo-3_*) or broadly in neurons (P*_rgef-1_*), plates with 20 synchronized L3-L4 larvae raised at 16°C were heat shocked at 34°C for 1 h followed by recovery at 20°C. After 24 h 20 animals were transferred to 0.5 mL M9 in 24-well plates and videos were immediately acquired at 20 frames per second on an Olympus SZX16 stereoscope equipped with a PLAPO 1x lens and an Olympus DP73 camera. Unbiased quantification of locomotion (expressed as body-bends per second) was performed using the wrMTrck plugin for ImageJ/Fiji (Nussbaum-Krammer *et al.* 2015). For strains with FLP D5 expressed in mechanosensory neurons (P*_mec-7_*) or serotonin producing neurons (P*_tph-1_*), synchronized L4 larvae were raised at 16°C and heat shocked at 33°C for 1 h followed by recovery at 20°C. Mechanosensation was evaluated after 24 h by gently touching animals laterally on their head 1-3 times with an eyelash mounted on a toothpick. Animals that did not respond immediately were scored as Mec (Mechanosensation defective). Rate of egg laying was determined during a 3 h time window 24 h after the heat shock.

## Results and Discussion

### Evaluation of FLP variants

Previous reports on FLP activity in *C. elegans* were mainly based on expression from extrachromosomal arrays containing typically hundreds of copies of the recombinase. To achieve completely stable meiotic and mitotic inheritance of the FLP transgene we decided to optimize FLP activity from single copy integrations into known sites of the *C. elegans* genome (MosSCI; (Frokjaer-Jensen *et al.* 2012)). We first designed a MosSCI dual color reporter integrated on chromosome II to enable direct visual scoring. The reporter contains the *hsp-16.41* promoter, which, in the absence of FLP activity, drives widespread and strong expression of mCherry (mCh) fused to the histone HIS-58 upon a short heat shock (Figure 1A; see also Figure 3). The mCh::*his-58* fusion gene is also regulated by the *let-858* 3’UTR, which serves as transcriptional terminator. The mCh::*his-58*::*let-858* 3’UTR cassette is flanked by two FLP recognition target (Frt) sequences, which trigger excision of the cassette in the presence of the FLP enzyme. Recombination between the two Frt sequences juxtaposes the *hsp-16.41* promoter with a GFP::*his-58*::*unc-54* 3’UTR fusion gene, which upon heat shock produces bright green nuclei. Thanks to the transparency of *C. elegans*, the number of green nuclei can be counted in live, anaesthetized animals on a compound microscope, providing a measure on the efficiency of FLP activity. We never observed GFP::HIS-58 expression in the absence of FLP, which indicates that the *let-858* 3’UTR sequence efficiently prevents transcriptional read-through (Davis *et al.* 2008; Voutev and Hubbard 2008). However, one report based on a multi-copy extrachromosomal array has suggested that a gene placed downstream of the *let-858* 3’UTR might be expressed at low levels (Ruaud *et al.* 2011). Thus, depending on the particular experiment it is advisable to investigate potential effects caused by the downstream gene in the absence of recombination.

Recombination efficiency can be expressed either as the percentage of animals expressing the downstream gene in at least a single cell or as the frequency of nuclei in which recombination has occurred. The former is the most feasible measurement for tissues with a high number of nuclei, whereas the latter is a more precise predictor of how many cells of a particular cell type will be affected in a given experiment (Table 1). Importantly, it should be noted that we might falsely score nuclei as GFP negative if the induction of the *hsp-16.41* promoter is weak, particularly if these nuclei are positioned deeper into the animal relative to the microscope objective. Thus, FLP activity might be underestimated for some tissues.

**Table 1.**
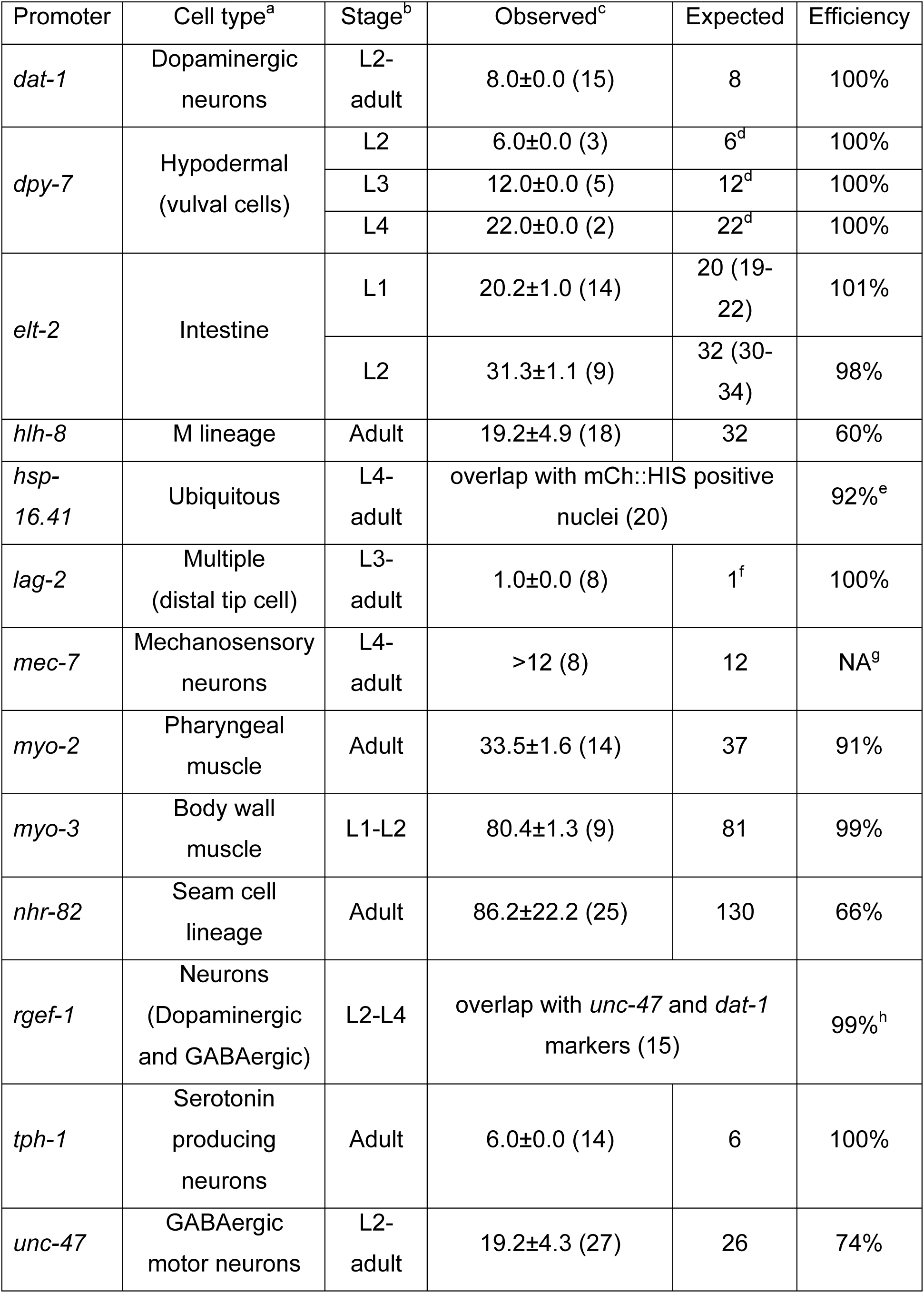
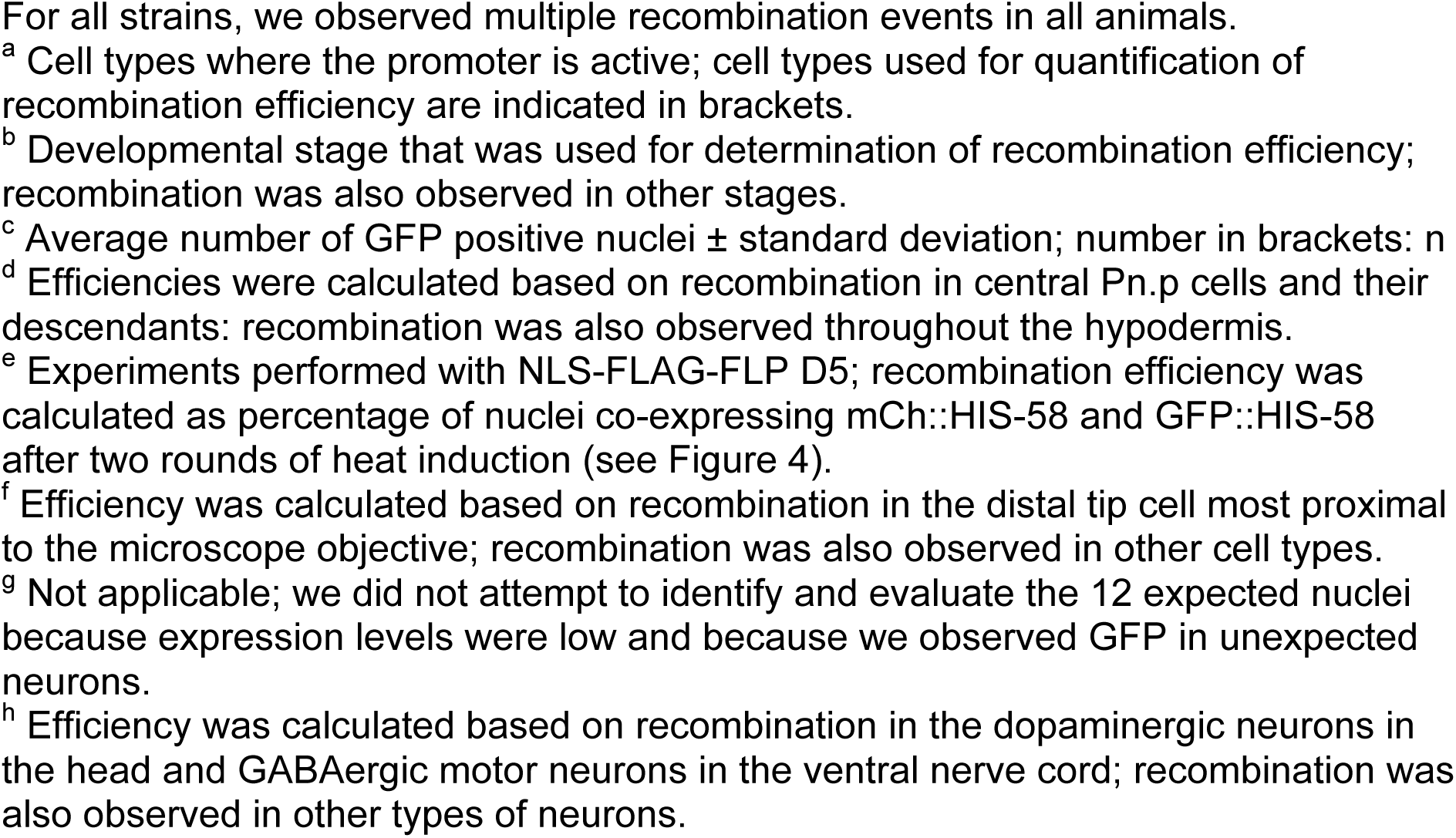
Quantification of FLP D5 activity in different tissues using the dual color reporter (see Figure 1).

| Promoter | Cell type<sup>a</sup> | Stage<sup>b</sup> | Observed<sup>c</sup> | Expected | Efficiency |
| --- | --- | --- | --- | --- | --- |
| *dat-1* | Dopaminergic neurons | L2-adult | 8.0±0.0 (15) | 8 | 100% |
| *dpy-7* | Hypodermal (vulval cells) | L2 | 6.0±0.0 (3) | 6<sup>d</sup> | 100% |
|  |  | L3 | 12.0±0.0 (5) | 12<sup>d</sup> | 100% |
|  |  | L4 | 22.0±0.0 (2) | 22<sup>d</sup> | 100% |
| *elt-2* | Intestine | L1 | 20.2±1.0 (14) | 20 (19-22) | 101% |
|  |  | L2 | 31.3±1.1 (9) | 32 (30-34) | 98% |
| *hlh-8* | M lineage | Adult | 19.2±4.9 (18) | 32 | 60% |
| *hsp-16.41* | Ubiquitous | L4-adult | overlap with mCh::HIS positive nuclei (20) |  | 92%<sup>e</sup> |
| *lag-2* | Multiple (distal tip cell) | L3-adult | 1.0±0.0 (8) | 1<sup>f</sup> | 100% |
| *mec-7* | Mechanosensory neurons | L4-adult | >12 (8) | 12 | NA<sup>g</sup> |
| *myo-2* | Pharyngeal muscle | Adult | 33.5±1.6 (14) | 37 | 91% |
| *myo-3* | Body wall muscle | L1-L2 | 80.4±1.3 (9) | 81 | 99% |
| *nhr-82* | Seam cell lineage | Adult | 86.2±22.2 (25) | 130 | 66% |
| *rgef-1* | Neurons (Dopaminergic and GABAergic) | L2-L4 | overlap with *unc-47* and *dat-1* markers (15) |  | 99%<sup>h</sup> |
| *tph-1* | Serotonin producing neurons | Adult | 6.0±0.0 (14) | 6 | 100% |
| *unc-47* | GABAergic motor neurons | L2-adult | 19.2±4.3 (27) | 26 | 74% |

For all strains, we observed multiple recombination events in all animals.
<sup>a</sup> Cell types where the promoter is active; cell types used for quantification of recombination efficiency are indicated in brackets.
<sup>b</sup> Developmental stage that was used for determination of recombination efficiency; recombination was also observed in other stages.
<sup>c</sup> Average number of GFP positive nuclei ± standard deviation; number in brackets: n
<sup>d</sup> Efficiencies were calculated based on recombination in central Pn.p cells and their descendants: recombination was also observed throughout the hypodermis.
<sup>e</sup> Experiments performed with NLS-FLAG-FLP D5; recombination efficiency was calculated as percentage of nuclei co-expressing mCh::HIS-58 and GFP::HIS-58 after two rounds of heat induction (see Figure 4).
<sup>f</sup> Efficiency was calculated based on recombination in the distal tip cell most proximal to the microscope objective; recombination was also observed in other cell types.
<sup>g</sup> Not applicable; we did not attempt to identify and evaluate the 12 expected nuclei because expression levels were low and because we observed GFP in unexpected neurons.
<sup>h</sup> Efficiency was calculated based on recombination in the dopaminergic neurons in the head and GABAergic motor neurons in the ventral nerve cord; recombination was also observed in other types of neurons.

We assumed that MosSCI FLP transgenes would be expressed at lower levels than transgenes on multicopy arrays so we engineered a novel FLP-encoding sequence with optimized codon usage for *C. elegans* and artificial introns ((Redemann *et al.* 2011); see Materials and Methods). Using the *nhr-82* promoter and the *glh-2* 3’UTR to control expression of FLP containing a glycine residue at position 5 (FLP G5; (Davis *et al.* 2008)) we first compared the activities of extrachromosomal and MosSCI lines on the dual color reporter. *nhr-82* is expressed in the seam cell lineage, which gives rise to a total of 130 seam and hypodermal nuclei in adults (Miyabayashi *et al.* 1999). We found that animals expressing FLP G5 from an extrachromosomal array had 99±21 GFP positive nuclei (average ± standard deviation; n=7), whereas animals expressing the same FLP version from an integrated single-copy had 40±20 GFP positive nuclei (n=11; Figure 2A).

**Figure 2.**
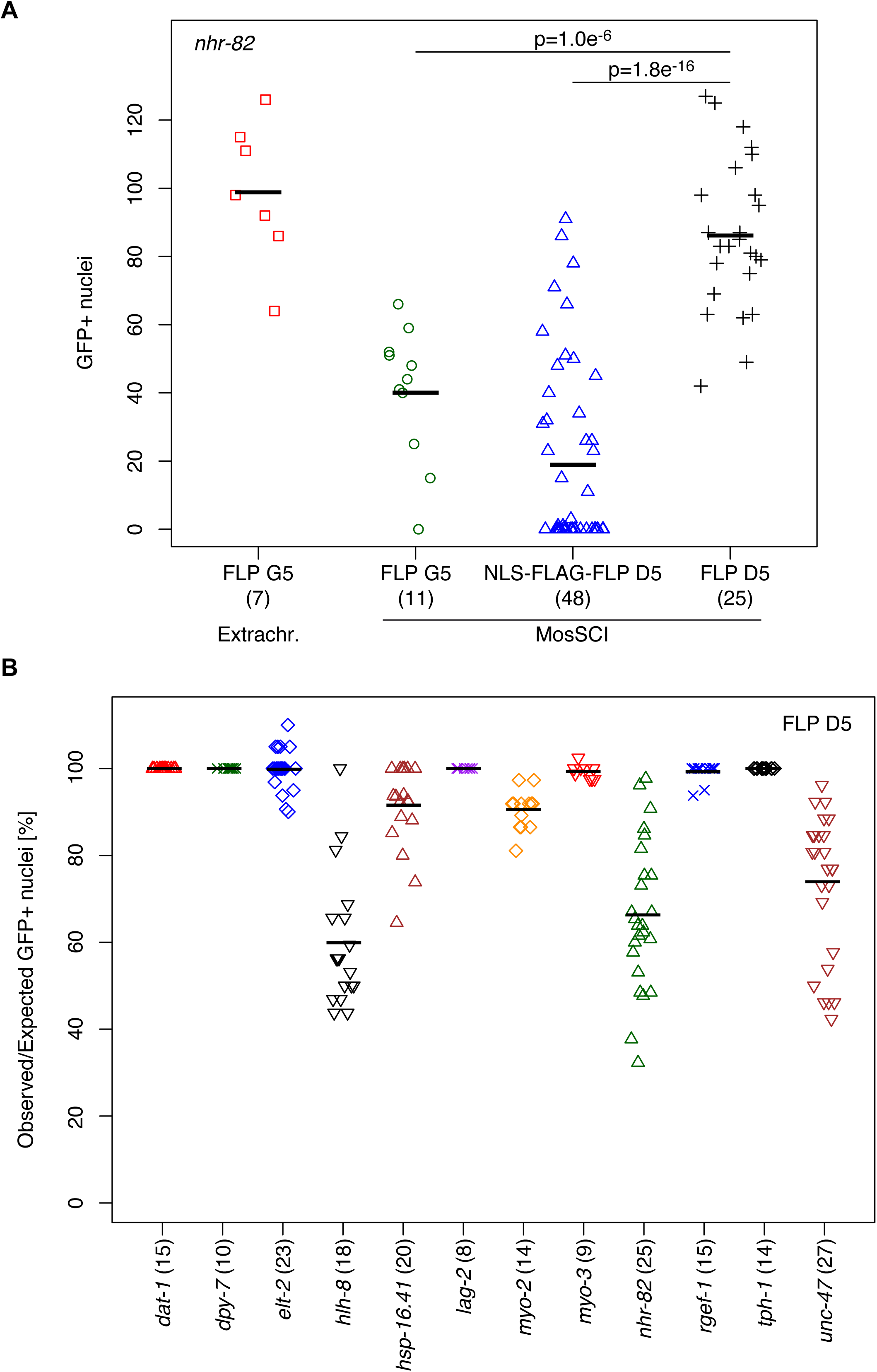
Optimization and quantification of FLP activity. FLP activity was evaluated based on the number of nuclei expressing GFP::HIS-58 in animals carrying the integrated dual color reporter described in Figure 1. (A) Comparison of strains expressing FLP from an extrachromosomal array (FLP G5; red squares) or from integrated single-copy transgenes (MosSCI): FLP G5 (green circles), NLS-FLAG-FLP D5 (blue triangles) or FLP D5 (black crosses). The same *nhr-82* promoter sequence was used in all strains. Shown are absolute numbers of GFP positive nuclei; numbers in brackets indicate the number of adults evaluated for each strain; black horizontal lines represent the means. P values were obtained through unpaired, two-sided t-tests, indicating that FLP D5 has the highest activity among the MosSCI strains. (B) Comparison of MosSCI strains expressing FLP D5 under control of different cell type-specific or inducible promoters: dopaminergic neurons (*dat-1*; red triangles), hypodermis (*dpy-7*; green crosses), intestine (*elt-2*; blue diamonds), M lineage (*hlh-8*; black triangles), heat inducible (*hsp-16.41*; brown triangles), notch signaling (*lag-2*; purple crosses), pharynx (*myo-2*; orange diamonds), body wall muscles (*myo-3*; red triangles), seam cell lineage (*nhr-82*; green triangles), pan-neuronal (*rgef-1*; blue crosses), serotonin producing neurons (*tph-1*; black diamonds), and GABAergic motor neurons (*unc-47*; brown triangles). Shown are the percentages of expected nuclei expressing GFP::HIS-58; numbers in brackets indicate the number of animals evaluated for each strain; black horizontal lines represent the means. For three promoters, on average, FLP activity was observed in 60-70% of the expected nuclei; for the remaining promoters the mean activity was >90% (see Table 1).

**Figure 3.**
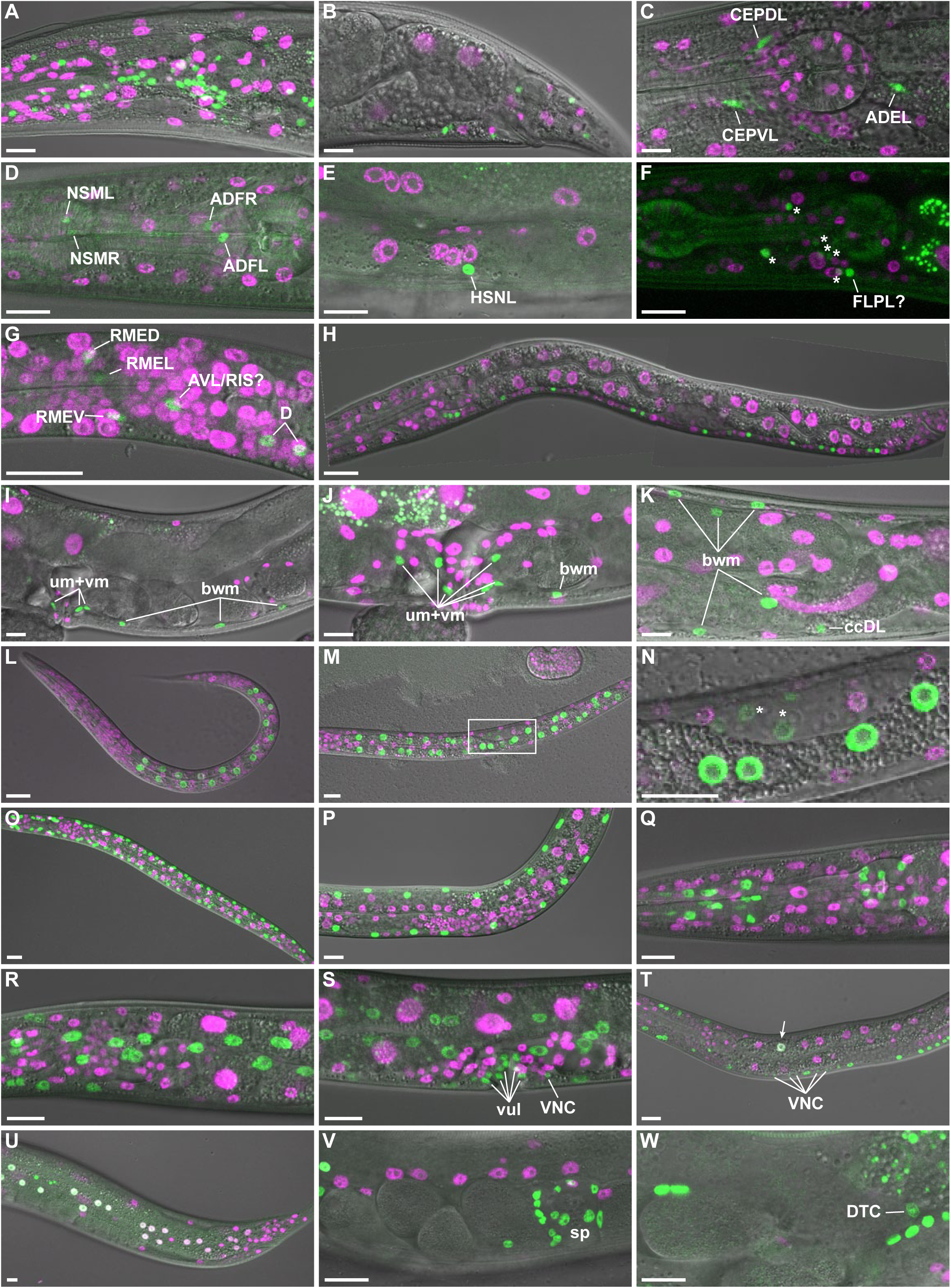
Tissue-specific FLPase activity. Transgenic animals expressing FLP D5 in specific cell types and the dual color reporter were observed by live confocal microscopy. (A-B) Promoter of *rgef-1* (P*_rgef-1_*); projection of eight confocal sections (A; adult head lateral view) or single confocal section (B; adult tail lateral view). (C) P*_dat-1_*; projection of four confocal sections of head of L4 larva. Note that the green channel represents co-expressed GFP::HIS-58 and mNeonGreen in this strain (see Supplementary Figure 3). (D-E) P*_tph-1_*; projection of three confocal sections (D; adult head lateral view) or a single confocal section (E; adult vulva lateral view). (F) P*_mec-7_*;projection of seven confocal sections of head of L4 larva. To ease visualization of the weak GFP signal the corresponding DIC image was not included in the merge. Asterisks indicate six unidentified neurons, whereas a seventh neuron might be FLPL. (G-H) P*_unc-47_*; late L1; projection of four confocal sections (G; head lateral view) or stitch of single confocal sections (H). (I-K) P*_hlh-8_*; single confocal section (I; adult vulva lateral view) or max projections (J; vulva region; eight confocal sections, K; posterior gonad loop region; fifteen confocal sections). Examples of body wall muscle (bwm), coelomocyte (cc), uterine muscle (um) and vulval muscle (vm) cells are indicated. (L-N) P*_elt-2_*; L1 (L; projection of seven confocal sections) and L2 larvae (M; projection of seven confocal sections, N; single confocal section corresponding to boxed area in M; note weak GFP expression in the gonad primordium [*]). (O-P) P*_myo-3_*; projection of seven confocal sections of L2 (O) and L4 (P) larvae. (Q) P*_myo-2_*; projection of twelve confocal sections of adult head. (R-T) P*_dpy-7_*; projection of eight (R; head lateral view) or ten (S; vulva lateral view) confocal sections of L4 larva or single confocal section of L3 larva (T). Examples of vulva cells (vul) and ventral nerve cord neurons (VNC) are indicated. Arrow points to intestinal cell that expresses both markers. (U) P*_nhr-82_*; projection of four confocal sections of tail of young adult. Note that white nuclei indicate simultaneous expression of mCh::HIS-58 and GFP::HIS-58. (V-W). P*_lag-2_*; projection of six confocal sections (V) or single confocal section (W) of central region of young adult. Spermatheca (sp) and distal tip cell (DTC) are indicated. All micrographs are oriented with anterior to the left and ventral down. Scale bar 10 μm.

The relative low efficiency of the MosSCI FLP G5 line led us to test a FLP variant with an aspartic acid residue at position 5, which was reported to be much more active in *Drosophila melanogaster* (FLP D5; (Nern *et al.* 2011)). Moreover, inspired by modifications of the Cre recombinase reported by (Ruijtenberg and van den Heuvel 2015) we also generated a FLP D5 variant with a heterologous nuclear localization signal and a 3xFLAG epitope, both at the N-terminus (FLAG-NLS-FLP D5). Surprisingly, animals expressing FLAG-NLS-FLP D5 had a highly variable number of GFP positive nuclei with many animals showing no recombination (19±27; n=48), whereas untagged FLP D5 was much more active (86±22 GFP positive nuclei; n=25), equaling the multicopy FLP G5 array (p=0.19 by unpaired, two-sided t-test; Figure 2A). It should be noted that in all experiments, animals were maintained at 16°C to minimize transcription from the *hsp-16.41* promoter of our reporters prior to heat-induction and recovery. Because FLP has a temperature optimum of 30°C (Buchholz *et al.* 1996), we speculated that higher recombination efficiencies could be obtained if animals were raised at 25°C. However, we obtained similar efficiencies at 16°C and 25°C (Supplementary Figure 1). Importantly, we only observed GFP positive nuclei in seam cells and the hypodermis, suggesting that the *nhr-82* promoter activates the FLP recombinase exclusively in the expected cell types. We therefore conclude that the combination of a *hsp-16.41*-controlled Frt construct and expression of FLP D5 in the seam cell lineage from integrated single-copy transgenes provides efficient spatiotemporal control of gene expression.

### Generation of tissue-specific FLP lines

We next generated multiple MosSCI lines expressing FLP D5 in specific cell types. In particular, to demonstrate the versatility of the FLP/Frt system we were interested in developing tools to control expression either broadly in the entire nervous system or restricted to specific neuronal sub-types. As pan-neuronal promoters we tested *unc-119* and *rgef-1*. Although expression of FLP D5 from the *unc-119* promoter induced recombination in neurons we also detected several green non-neuronal nuclei (Supplementary Figure 2A). The activity of *unc-119* outside the nervous system has been reported in WormBase (http://www.wormbase.org) and we did not characterize this construct further. Instead we focused on *rgef-1*, which is expressed in post-mitotic neurons (Altun-Gultekin *et al.* 2001). In all P*_rgef-1_*::FLP D5 animals (n>40), we observed numerous green nuclei throughout the nervous system (Figure 3A-C; note that several of the images represent single confocal sections, hence only subsets of nuclei are visible; Table 1). We did not attempt to count the absolute number of GFP positive nuclei, but when crossed to markers of dopaminergic and GABAergic motor neurons, we observed recombination in 99% of the cells expressing the marker (n=15; Table 1; Supplementary Figure 2C-D). Thus, we conclude that the *rgef-1* promoter is highly active and specific for induction of recombination in *C. elegans* neurons. As examples of specific neuronal subtypes, we tested four promoters: *dat-1* (dopaminergic neurons; (Nass *et al.* 2002)), *mec-7* mechanosensory and other neurons; (Mitani *et al.* 1993)), *tph-1* (serotonin producing neurons; (Sze *et al.* 2000)) and *unc-47* (GABAergic motor neurons; (Mcintire *et al.* 1997)). In all 14 P*_tph-1_*::FLP D5 animals analyzed we observed GFP expression exclusively in six neurons that were located in the expected positions (NSML/R, ADFL/R and HSNL/R; Figures 2B and 3D-E; Table 1). For the *unc-47* promoter, we observed 17±5 GFP positive nuclei in the head and along the ventral nerve cord, which corresponds to 64% of the expected nuclei (Figures 2B and 3G-H; Table 1). Although the *nhr-82* and *unc-47* promoters produced similar recombination efficiencies in terms of percentage of GFP positive nuclei (Figure 2B; Table 1) they showed a remarkable difference: Most green nuclei in P*_nhr-82_* animals co-expressed mCh::HIS-58, which suggests that only one of the two alleles in the genome had undergone FLP-recombination (Figure 3U). In contrast, none of the green nuclei in P*_unc-47_* animals had expression of mCh::HIS-58, indicating that either both alleles had recombined or, less likely, only a single allele was expressed. Similar to the results with the *tph-1* and *unc-47* promoters, the P*_mec-7_*::FLP strain also produced recombination specifically in the nervous system, but in all 8 animals analyzed we observed more GFP positive nuclei than expected. For instance, we expected recombination only in two neurons in the head (FLPL/R) but we detected up to seven GFP positive nuclei (Figure 3F). Because the FLP/Frt system also tracks cell lineages (Voutev and Hubbard 2008), we speculate that these “extra” cells might reflect transient activation of the *mec-7* promoter earlier in development. An alternative, and not mutually exclusive, possibility is that current expression annotations are incomplete (Schmitt *et al.* 2012). The first line carrying a P*_dat-1_*::FLP D5 transgene did not produce recombination of the dual color reporter (data not shown), possibly due to insufficient expression of the recombinase. We therefore generated a second line, this time including a downstream transsplicing cassette encoding mNeonGreen to indirectly visualize FLP D5 expression. This line had robust mNeonGreen expression in six head neurons and two posterior neurons of all animals, as expected for the *dat-1* promoter (n=37; Supplementary Figure 3). When crossed with the dual color reporter, co-expression of GFP::HIS-58 and mNeonGreen was observed in all dopaminergic neurons (Figure 3C; Table 1; Supplementary Figure 3). Based on these results we conclude that the *C. elegans* nervous system is amenable to spatiotemporal control of gene expression using integrated single-copy FLP/Frt transgenes, in some cases with ~65%-100% efficiency.

The M lineage is derived from a single embryonic mesodermal blast cell, which generates 14 striated body wall muscle cells, 16 non-striated uterine and vulval muscle cells and 2 coelomocytes during larval development (Sulston and Horvitz 1977). Thus, the M lineage offers an attractive system to study postembryonic control of cell proliferation and differentiation. The *hlh-8* gene is a marker of the M lineage (Harfe *et al.* 1998) and we tested if the *hlh-8* promoter can be used to induce specific expression of FLP D5 and recombination in this cell lineage. In adults, we observed GFP expression in 19±5 nuclei, all belonging to the expected cell types (Figures 2B and 3I-K; Table 1). This demonstrates that the M lineage is susceptible to lineage-specific, FLP-mediated recombination via the *hlh-8* promoter. A similar conclusion was recently made based on the Cre/lox system although a low frequency of recombination outside the M lineage was also observed (Ruijtenberg and van den Heuvel 2015).

The *C. elegans* intestine consists of 20, or occasionally 19, 21 or 22 cells (Sulston and Horvitz 1977; Mcghee 2007). At the L1/L2 larval molt, 10-14 of the ~20 nuclei divide to produce a total of 30-34 nuclei: for calculation of recombination frequencies we assumed 20 and 32 nuclei in L1 and L2 larvae, respectively, as expected values. To drive expression of the recombinase in the intestine we cloned the *elt-2* promoter (Fukushige *et al.* 1998) and crossed P*elt-2*::FLP D5 animals with the dual color reporter strain. In L1 and L2 larvae we observed 20±1 and 31±1 GFP positive nuclei, respectively, which corresponds to 98-100% efficiency (Figures 2B and 3L-N; Table 1). We also found expression of GFP::HIS-58 in a few cells of the gonad primordium (asterisks in Figure 3N). Analyzing adult animals consistently revealed GFP positive nuclei in the somatic gonad, including the spermathecae and gonadal sheet cells (Supplementary Figure 2D). Activity of the *elt-2* promoter in the somatic gonad has been reported (Ruijtenberg and van den Heuvel 2015) and this should be taken into account when designing FLP or Cre-based recombination experiments to analyze biological processes in the intestine. Nevertheless, in L1 and young L2 larvae before proliferation of the gonad primordium the contribution from non-intestinal cells is likely to be negligible.

To achieve recombination in non-striated pharyngeal muscles and striated body wall muscles we tested the well-established *myo-2* and *myo-3* promoters (Okkema *et al.* 1993). Both promoters induced specific recombination in the expected cell types at 91-99% efficiency with 34±2 and 80±1 GFP positive nuclei in the pharynx and the body wall muscles, respectively (Figures 2B and 3O-Q; Table 1). As described above, using the *nhr-82* promoter to control FLP D5 expression leads to recombination in nuclei that will later form part of the hypodermis. However, many hypodermal nuclei are not derived from the seam cell lineage so to target the hypodermis more broadly we tested the *dpy-7* promoter (Gilleard *et al.* 1997; Page and Johnstone 2007). All animals analyzed (n>50) had very high density of GFP positive nuclei in the hypodermis, as well as in the ventral nerve cord and in vulval cells (Figure 3R-T; Table 1). The expression of GFP in these latter cell types suggests that P_*dpy-7*_::FLP D5-mediated recombination happens relatively early in development before the P cells migrate and produce neurons and vulva precursor cells (Sulston and Horvitz 1977). Moreover, we also observed partial recombination in a few polyploid intestinal cells in some animals (1.4±2.2 nuclei, n=50; arrow in Figure 3T). Finally, we included the *lag-2* promoter in our battery of FLP D5 lines to explore the potential of the FLP/Frt system in the context of a classical cell-signaling pathway. LAG-2 is a ligand for GLP-1 and LIN-12 Notch-like receptors and is expressed in many different cell types to influence proliferation and differentiation of neighboring cells (Henderson *et al.* 1994; Wilkinson *et al.* 1994; Greenwald 2005). In all P*_lag-2_*::FLP D5 animals (n>50) we observed a large number of GFP positive nuclei, in particular in the somatic gonads including the spermathecae and the distal tip cells (Figure 3V-W; Table 1).

Based on the results above, we conclude that single-copy FLP D5 transgenes are highly efficient and specific tools to excise Frt-flanked DNA fragments from the *C. elegans* genome. Because the FLP D5 transgenes are integrated into the genome they can easily be crossed to relevant Frt strains and show complete stability during meiotic and mitotic divisions, ensuring that all cells in individual animals and all animals in a population carry the FLP D5 transgene.

### Temporal control of leaky transcription

Some protocols utilize the basal or “leaky” transcription of the *hsp-16.41* promoter in non-induced conditions to achieve low expression levels. This is typically the case in DamID experiments where low amounts of a chromatin-interacting protein fused to *Escherichia coli* DNA adenine methyltransferase (Dam) are used to identify protein-DNA interactions (Gomez-saldivar *et al.* 2016). This low expression level is typically not detectable by fluorescent reporters or Western blotting but is sufficient to generate specific DamID signals (Figure 1C, 0 h post heat shock; see Supplementary Figure 1 in (Gonzalez-Aguilera *et al.* 2014)). However, because methylation marks made by Dam are stable in post-mitotic cells, DamID profiles obtained from old animals will represent the sum of methylation accumulated since their cells were born. Therefore, to determine protein-DNA interactions taking place in old animals expression of the Dam fusion protein should be completely avoided at earlier time points. To this end, we tested if the *hsp-16.41* promoter can be used to temporally control FLP-mediated recombination (Figure 4A). Using again the dual color reporter to evaluate recombination efficiency, we observed that an initial heat shock led to ubiquitous and strong expression of mCh::HIS-58, whereas similar GFP::HIS-58 expression was only observed after a second heat shock (Figure 4B). These results demonstrate that recombination requires induction of the *hsp-16.41* promoter to produce sufficient FLP enzyme and that once this occurs; recombination is efficient throughout the animal. We have implemented this system to analyze nuclear organization in old animals and preliminary data suggest both local and global changes when compared to young animals (Muñoz-Jiménez *et al*; unpublished observations). Moreover, by substituting the *hsp-16.41* promoter in the Frt construct with a tissue-specific promoter, the system can easily be adapted to stimulation within a target tissue using heat shock (Figure 1B). For instance, if a BirA fusion gene is inserted after the Frt-flanked cassette, the system can be used for spatiotemporal control of proximity-dependent biotin identification (BioID) of protein interaction partners (Kim and Roux 2016). Whether the best configuration to achieve spatiotemporal control is by expression of FLP from a tissue-specific promoter and the Frt construct from a heat-inducible promoter or vice versa depends on the particular experiment. In the former configuration, potentially the entire cell lineage(s) in which FLP has been transiently expressed will rapidly produce a boost of temporal expression of the downstream gene upon heat shock (Figure 1A, C). In contrast, if the recombinase is placed under control of the *hsp-16.42* promoter, a heat shock will induce expression of the downstream gene only in cells in which the tissue-specific promoter is currently active and with a time-delay reflecting the kinetics of recombination (Figure 1B, D). Typically, the expression levels in the latter case will be lower than the ones obtained with the induced *hsp-16.42* promoter, although this will naturally depend on the particular tissue-specific promoter.

**Figure 4.**
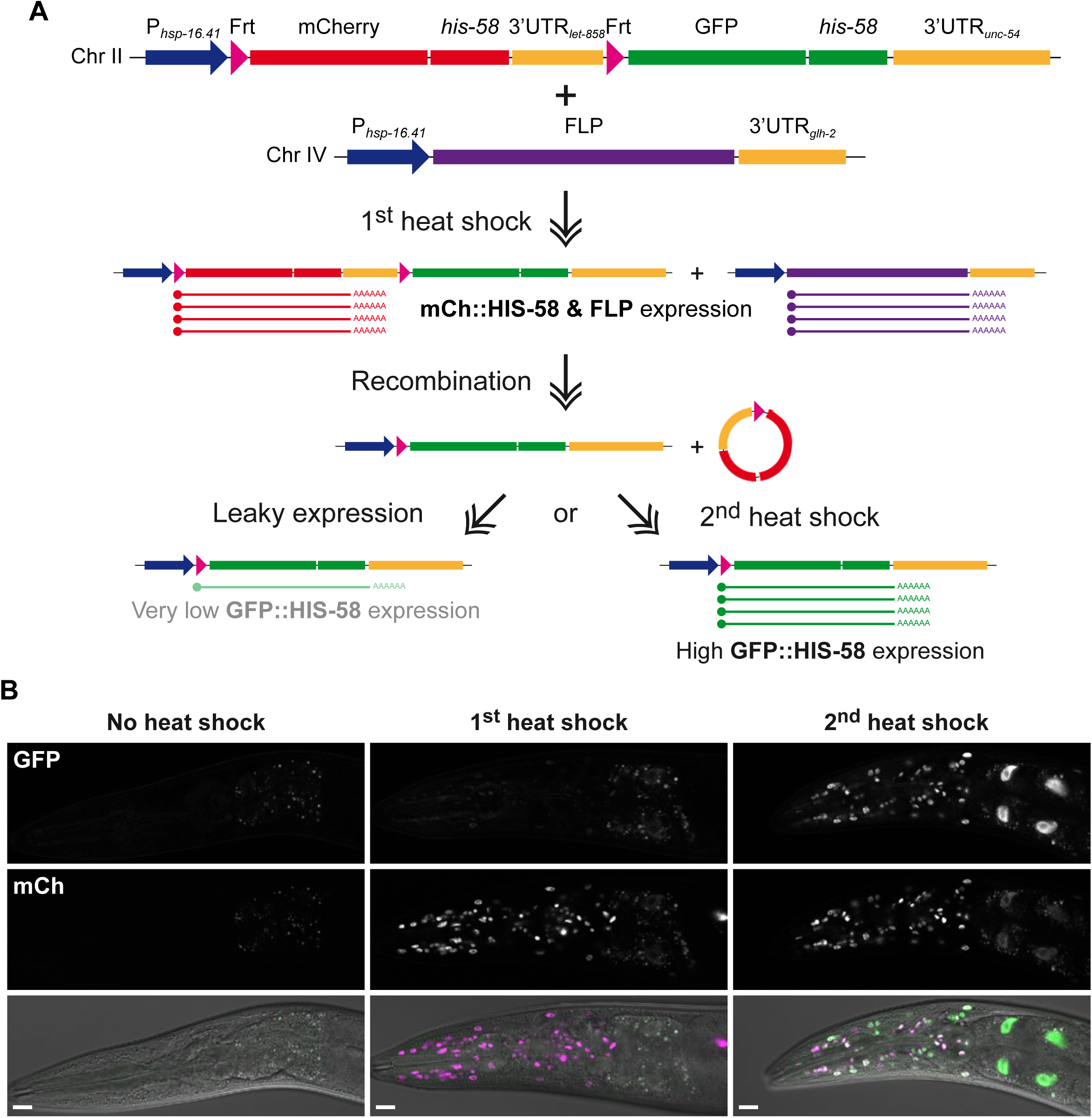
Temporal control of FLP recombination. Frt-flanked cassettes can be efficiently excised by heat shock-induced FLP expression. (A) Schematic representation of experimental setup and (B) single confocal sections of adult heads. The first time the animals are heat shocked they express mCh::HIS-58 and FLP. The FLP recombinase excises the mCh::*his-58* cassette but efficient expression of the downstream GFP::*his-58* fusion gene requires a second heat shock, suggesting that recombination is completed after the transient activation of the *hsp-16.41* promoter ends. Scale bar 10 μm.

### Spatiotemporal control of cell ablation

Killing of specific cells or cell types represents an important tool to understand their physiological role. Different systems have been proposed to achieve temporal and/or spatial control, including expression of reconstituted caspases (Chelur and Chalfie 2007), tetanus toxin (Davis *et al.* 2008), miniSOG (Qi *et al.* 2012; Xu and Chisholm 2016) and histamine-gated chloride channels (Pokala *et al.* 2014). Each of these methods has its advantages and drawbacks and we reasoned that our collection of stable FLP lines could potentially serve as an efficient and fast way to target a variety of specific tissues without the need of multiple cloning steps or experience with optogenetic equipment. To test this, we used a cDNA encoding the PEEL-1 toxin (Seidel *et al.* 2011) to replace the GFP::*his-58* sequence of our dual color reporter and inserted this construct as single copy transgene on chromosome II (Figure 5A). We first crossed the *peel-1* line to the P*_rgef-1_*::FLP D5 and P*_myo-3_*::FLP D5 lines to broadly target neurons and muscles, respectively. For unbiased assessment of locomotion we transferred the animals to liquid media and quantified the number of body-bends per second using the wrMTrck software (Nussbaum-Krammer *et al.* 2015). In the absence of heat shock all lines behaved like wild type animals, indicating that leaky expression of PEEL-1 in neurons or muscles has no obvious consequence (blue boxes in Figure 5B; Videos 1-2). In contrast, when PEEL-1 expression was induced in muscles, all animals became uncoordinated (Unc phenotype) and were paralyzed within 2 h (Video 3). Although the penetrance was less severe, *peel-1* P*_rgef-1_*::FLP D5 animals (Video 4) had a significant Unc phenotype, whereas the heat shock did not affect *peel-1* animals in the absence of FLP recombinase (red boxes in Figure 5B).

**Figure 5.**
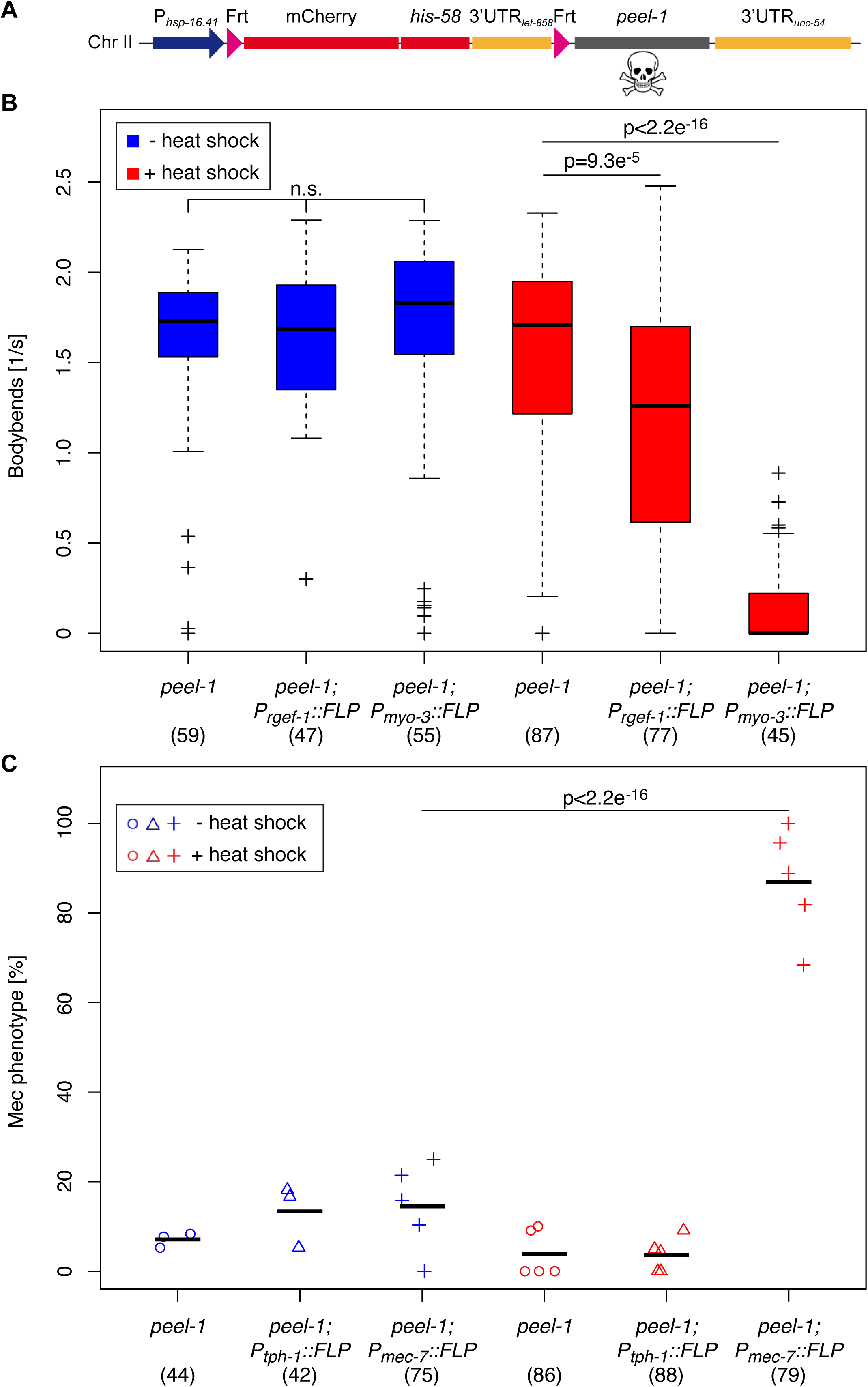
Spatiotemporal control of cell ablation. (A) Schematic representation of FLP-regulated *peel-1* cell killing construct. Heat shock activation of the construct produces mCh::HIS-58 in the absence of FLP and PEEL-1 toxin in cells positive for FLP activity. (B) Locomotion of animals carrying the *peel-1* construct alone or together with either P*_rgef-1_*::FLP or P*_myo-3_*::FLP was quantified before (blue boxes) or after heat shock (red boxes). Top and bottom of boxes indicate the 1^st^ and 3rd quartiles, respectively, whereas lines inside boxes correspond to the medians; whiskers cover the 1.5x interquartile range. (C) Strains carrying the *peel-1* construct alone (circles) or together with either P*_tph-1_*::FLP (triangles) or P*_mec-7_*::FLP (crosses) were observed for mechanosensation defects (Mec) in the absence (blue symbols) or after heat shock (red symbols). Shown are the percentages of Unc (B) and Mec (C) animals in 3-9 independent experiments; numbers in brackets indicate the total number of animals evaluated for each condition; black horizontal lines represent the means. P values were obtained through Mann-Whitney-Wilcoxon (B) and Fisher’s exact (C) tests, indicating that heat-induced ablation of neurons (P*_rgef-1_*::FLP), body wall muscles (P*_myo-3_*::FLP) or mechanosensory neurons (P*_mec-7_*::FLP) induce Unc or Mec phenotypes.

Next, we tested the cell ablation system in specific neuronal subtypes using the *tph-1* and *mec-7* promoters. As described above, *mec-7* is expressed in mechanosensory neurons and *mec-7* mutants have mechanosensation (Mec) phenotypes (Mitani *et al.* 1993). In concordance with this, we observed 87±12% Mec animals in heat-shocked *peel-1* P*_mec-7_*::FLP D5 animals (n=79), whereas FLP expression in serotonin producing neurons (from the *tph-1* promoter) did not affect mechanosensation (red symbols Figure 5C). As expected, in the absence of heat shock, the expression of the recombinase in any of the two neuronal subtypes did not produce Mec phenotypes (p=0.25-0.48 by Fisher’s exact test (blue symbols in Figure 5C). Mutation of *tph-1* interferes with egg laying (Sze *et al.* 2000) and *peel-1* P*_tph-1_*::FLP D5 animals produced fewer eggs per hour than *peel-1* animals after heat shock (p=0.02 by unpaired, two-sided t-test; Supplementary Figure 4). However, the transgene P*_tph-1_*::FLP D5 reduced egg laying also without heat shock (p=0.003), which could reflect that the serotonin producing neurons are potentially affected already by the leaky expression of PEEL-1 from the *hsp-16.41* promoter (Supplementary Figure 4). In conclusion, the tools developed here to induce killing of cells in a spatiotemporal manner produce in most cases the expected behavioral phenotypes. More detailed microscopy studies are required to determine the precise damage at tissue level and future experiments under comparable conditions are needed to determine which effector (e.g. PEEL-1, tetanus toxin, caspases, histamine-gated chloride channels, etc.) might be most suitable for recombination-driven cell ablation.

### Tissue-specific gene disruption by combined FLP and CRISPR tools

The development of CRISPR/Cas9 methods to precisely engineer the genome has revolutionized genetics experiments in a wide range of organisms, including *C. elegans* (Dickinson and Goldstein 2016). We decided to explore the combination of CRISPR and FLP-based recombination to interfere with gene expression in specific cells. As a proof of principle, we modified the endogenous *baf-1* locus. BAF-1 is a small chromatin-binding protein essential for nuclear envelope assembly (Gorjanacz *et al.* 2007). Homozygous *baf-1* mutants complete embryogenesis thanks to maternally contributed protein but arrest as larvae with defects in multiple tissues, whereas BAF-1-depleted embryos die during early embryogenesis (Gorjanacz *et al.* 2007; Margalit *et al.* 2007). We first generated a SapTrap compatible (Schwartz and Jorgensen 2016) GFP knockout (KO) cassette containing Frt sites in introns 1 and 2 (indicated as *g>f>p* in genotype descriptions). The two Frt sites are separated by 205 bp, which should be compatible with optimal recombination efficiency (Ringrose *et al.* 1999). When expressed in *C. elegans* cells without FLP activity the introns are removed from the primary transcript by splicing and the processed mRNA will encode GFP::BAF-1 that localizes to the nuclear envelope (Figure 6A; “Other tissues”). In contrast, in the presence of FLP the second GFP exon is excised by recombination between the two Frt sites, which cause a reading frame shift in the third GFP exon and the appearance of a premature termination codon (PTC; Figure 6A; “Intestine”). This will most likely lead to degradation of the aberrant GFP::*baf-1* transcript by nonsense-mediated mRNA decay, which is highly efficient in *C. elegans* (Zahler 2012). Even if a few mRNA molecules escape decay, expression of *baf-1* downstream of the PTC would require that ribosomes are not released at the PTC but instead re-initiate translation (He and Jacobson 2015).

**Figure 6.**
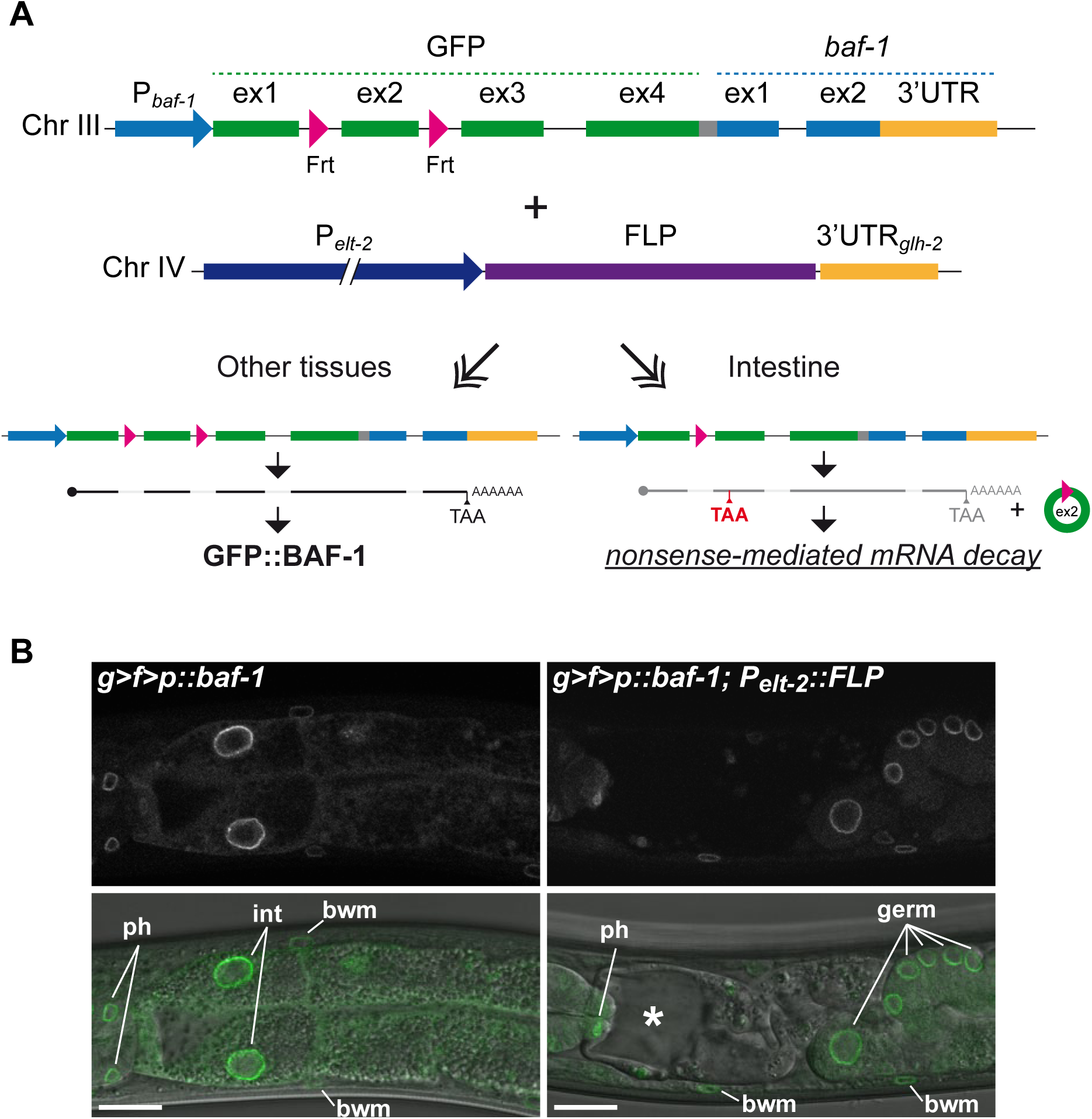
Tissue-specific inhibition of gene expression. (A) Schematic representation of experimental design and (B) single confocal sections of anterior part of intestine in L4 larvae. The endogenous *baf-1* locus was modified by insertion of a GFP knockout (KO) cassette immediately upstream of the coding region. The GFP KO cassette contains Frt sites in the first and second intron (indicated as *g>f>p* in B): these sites do not interfere with pre-mRNA splicing and, in the absence of FLP, the locus expresses GFP::BAF-1. In cells expressing FLP, GFP exon 2 is excised and the spliced mRNA contains a premature termination codon (red TAA), which triggers nonsense-mediated mRNA decay. (B) The left animal expresses GFP::BAF-1 in all cells, whereas the right animal lack expression in intestinal cells due to the specific expression of FLP in this tissue. Nuclei of different tissues are indicated: ph: pharynx; int: intestine; bwm: body wall muscle nuclei; germ: germ line. An asterisk indicates an enlarged intestinal lumen surrounded by abnormally thin cellular processes. Scale bar 10 μm.

Animals homozygous for the GFP KO cassette in the *baf-1* locus did not show any obvious phenotypes and bright fluorescent signal was detected at the periphery of nuclei throughout the body at all developmental stages (Figure 6B left and data not shown). However, when crossed with the P*_elt-2_*::FLP D5 line, the intestine showed clear signs of atrophy and lack of GFP expression, whereas other tissues as expected expressed GFP::BAF-1 (Figure 6B right). Concordantly, animals were sick with reduced body size and slow movements. A more detailed characterization of the role of BAF-1 in the development and/or maintenance of the intestine goes beyond the scope of this study but our data clearly demonstrate the potential to use this new collection of FLP lines to address gene function in specific tissues. We have focused on an example with insertion of the GFP KO cassette into an endogenous locus but the same strategy can easily be adapted for generation of Frt-flanked transgenes for Mos1-mediated single copy insertion into the genome as has recently been demonstrated for the Cre/lox system (Flavell *et al.* 2013; Ruijtenberg and van den Heuvel 2015). In the example presented here, we inserted the GFP KO cassette at the translational start codon and, as explained above, this is likely to efficiently prevent synthesis of the tagged proteins in FLP-expressing cells. We are currently testing the efficiency of conditionally controlling expression of loci in which the GFP KO cassette, or a novel red variant (wrmScarlet (El Mouridi *et al.* 2017) KO cassette is inserted immediately in front of the endogenous translational termination codon. We expect that the mRNA produced in FLP-expressing cells will be unstable due to the presence of a premature termination codon specifically in these cells, but this needs to be tested empirically.

## Conclusion

We have provided quantitative evidence that stable FLP lines can drive specific and efficient recombination between Frt sites in multiple cell types in *C. elegans*. Because the FLP transgenes are integrated into a known locus in the genome, genetic crosses to different Frt-flanked constructs are straightforward. We plan to expand the collection of FLP lines in the future and we hope other members of the community will participate to increase the number of specific cell types amenable to FLP-mediated recombination. Site-specific recombination can be utilized in many experimental strategies (see (Hubbard 2014) for a comprehensive review) and the FLP lines reported here can easily be combined with existing tools for tissue-specific GFP tagging (Schwartz and Jorgensen 2016) or expression of RNAi hairpins (Voutev and Hubbard 2008). We have demonstrated that the FLP system can be used tissue-specifically to knock out gene function or ablate cells. Our GFP knockout cassette is compatible with the efficient SapTrap protocol for preparation of repair templates for genome engineering (Schwartz and Jorgensen 2016) and can therefore readily be applied to other loci. Finally, we envision that other advances in CRISPR technology will further facilitate insertion of Frt sites into endogenous loci to make them controllable by FLP.

## Acknowledgment

We are grateful to Alejandro Ferreiro Morales, Carmen Serrano Tasset, Cristina González-Aguilera, Eric Jorgensen and Esther Dalfo for plasmids as well as to Nuria Flames for suggestions on neuron-specific promoters and to Anita Fernandez for critical reading of the manuscript. We also wish to acknowledge Addgene for sequence verifications, the Caenorhabditis Genetics Center (CGC), which is funded by NIH Office of Research Infrastructure Programs (P40 OD010440) and WormBase, which is supported by a grant from the National Human Genome Research Institute at the NIH # U41 HG002223 and the British Medical Research Council. This project was funded by the Spanish Ministry of Economy and Competitiveness (BFU2013- 42709P and BFU2016-79313-P) and the European Regional Development Fund.

## Figure legends

Supplementary Figure 1. Efficiency of FLP-mediated recombination is temperature-independent. Transgenic animals expressing FLP in GABAergic motor neurons from the *unc-47* promoter were raised at either 16°C or 25°C prior to heat induction of the dual color reporter. The number of GFP positive nuclei was observed by live confocal microscopy and represented as percentage of expected nuclei. Numbers in parentheses indicate sample sizes; black horizontal lines represent the medians. P value was obtained through an unpaired, two-sided t-test, indicating that recombination efficiency is temperature independent.

Supplementary Figure 2. Evaluation of specificity and efficiency of FLP-mediated recombination. Transgenic animals expressing FLP in specific cell types and the dual color reporter were observed by live confocal microscopy. (A) Maximum projections of confocal micrographs of heads of adults expressing FLP G5 under control of the *unc-119* promoter from either an extrachromosomal array (left panels) or an integrated single-copy transgene (right panels). The frequency of recombination as visualized by the number of GFP positive nuclei is very low in the MosSCI strain, whereas many neurons and non-neuronal nuclei express GFP::HIS-58 in the array strain. Nuclei in which recombination has not taken place express mCherry::HIS-58 (mCh; magenta in merge). (B) Maximum projections of confocal micrographs of central part of adults expressing FLP D5 under control of the *elt-2* promoter from an integrated single-copy transgene. In addition to expression in the large intestinal nuclei as expected, GFP::HIS-58 is also observed in gonadal sheath cells (gs; left panels), the spermathecae (sp) and uterine cells (u; right panels). (C) Single confocal section of central part of L4 larva expressing FLP D5 under control of the *rgef-1* promoter from an integrated single-copy transgene as well as diffuse mCherry from the *unc-47* promoter to mark GABAergic motor neurons. White arrows point to nuclei of GABAergic motor neurons, which all have undergone recombination and express GFP::HIS-58. (D) Maximum projections of three confocal micrographs of head of L4 larva expressing FLP D5 under control of the *rgef-1* promoter from an integrated single-copy transgene as well as diffuse mCherry from the *dat-1* promoter to mark dopaminergic neurons. White arrows point to nuclei of dopaminergic neurons, which all have undergone recombination and express GFP::HIS-58. Scale bar 10 μm.

Supplementary Figure 3. Evaluation of specificity and efficiency of FLP-mediated recombination in dopaminergic neurons. Single confocal sections of live animals co-expressing FLP D5 and mNeonGreen from a bicistronic transgene under control of the *dat-1* promoter either in the absence or the presence of the dual color reporter (left and right micrographs, respectively). Shown are examples of CEP (A) and PDE (B) neurons. In the merged images GFP+mNeonGreen and mCherry are represented in green and magenta, respectively; in the individual inserts GFP+mNeonGreen and mCherry are represented with Fiji’s ‘Fire’ and ‘Gray’ lookup tables, respectively. The brighter GFP+mNeonGreen signal and the absence of mCherry signal in the dopaminergic neurons (arrows in right micrographs) demonstrate the specificity and efficiency of recombination. Scale bar 10 μm.

Supplementary Figure 4. Rate of egg laying after induced cell ablation. Three strains were evaluated: one that carries the P*_hsp-16.41_*>*mCh::his-58*>*peel-1* construct (see Figure 5A) alone and two that carry the *peel-1* construct together with a P*_tph-1_*::FLP D5 or a P*_mec-7_*::FLP D5 transgene. Two to four adults where incubated together for three hours and the average rate of egg laying per animal per hour was determined. A total of 12 plates for each condition were evaluated; numbers in brackets indicate total number of animals analyzed. Blue symbols report the observations from control animals, whereas red symbols correspond to animals evaluated 24-27 h after 15 min heat shock. Animals expressing FLP in serotonergic neurons (P*_tph-1_*::FLP) laid fewer eggs under both conditions. P values were obtained through unpaired, two-sided t-tests.

Video S1. Transgenic P*_hsp-16.41_*>*mCh::his-58*>*peel-1*; P*_myo-3_*::FLP D5 animals observed without heat shock. Scale bar corresponds to ~120 μm.

Video S2. Transgenic P*_hsp-16.41_*>*mCh::his-58*>*peel-1*; P*_rgef-1_*::FLP D5 animals observed without heat shock. Scale bar corresponds to ~120 μm.

Video S3. Transgenic P*_hsp-16.41_*>*mCh::his-58*>*peel-1*; P*_myo-3_*_>_::FLP D5 animals observed 6 h after 15 min heat shock. Scale bar corresponds to ~120 μm.

Video S4 Transgenic P*_hsp-16.41_*>*mCh::his-58*>*peel-1*; P*rgef-1*::FLP D5 animals observed 6 h after 15 min heat shock. Scale bar corresponds to ~120 μm.

